# Risk factor analysis of equine strongyle resistance to anthelmintics

**DOI:** 10.1101/158105

**Authors:** G. Sallé, J. Cortet, I. Bois, C. Dubès, Q. Guyot-Sionest, C. Larrieu, V. Landrin, G. Majorel, S. Wittreck, E. Woringer, A. Couroucé, J. Guillot, P. Jacquiet, F. Guégnard, A. Blanchard, A. Leblond

## Abstract

Intestinal strongyles are the most problematic endoparasites of equids as a result of their wide distribution and the spread of resistant isolates throughout the world. While abundant literature can be found on the extent of anthelmintic resistance across continents, empirical knowledge about associated risk factors is missing. This study brought together results from anthelmintic efficacy testing and risk factor analysis to provide evidence-based guidelines in the field. It involved 688 horses from 39 French horse farms and riding schools to both estimate Faecal Egg Count Reduction (FECR) after anthelmintic treatment and to interview farm and riding school managers about their practices. Risk factors associated with reduced anthelmintic efficacy in equine strongyles were estimated across drugs using a marginal modelling approach. Results demonstrated ivermectin efficacy (96.3% FECR), the inefficacy of fenbendazole (42.8% FECR) and an intermediate profile for pyrantel (90.3% FECR). Risk factor analysis provided support to advocate for FEC-based treatment regimens combined with individual anthelmintic dosage and the enforcement of tighter biosecurity around horse introduction that contributed to lower drug resistance risk by 1.75. Premises falling under this typology also relied more on their veterinarians suggesting they play an important role in the sustainability of anthelmintic usage. Similarly, drug resistance risk was halved in premises with frequent pasture rotation and with stocking rate below five horses/ha. This is the first empirical risk factor analysis for anthelmintic resistance in equids, whose findings should guide the implementation of more sustained strongyle management in the field.

## 1. Introduction

The diversity of helminth species infecting horses is large, and differences in life cycles, epidemiology, pathogenicity and drug susceptibility make it increasingly challenging to define good sustainable parasite control programs. Strongyles remain a major concern. They can be classified into two sub-families, namely Strongylinae (large strongyles) and Cyathostominae known as small strongyles or cyathostomins (Lichtenfels et al., 2008). The large strongyle *Strongylus vulgaris* is associated with a high mortality rate resulting from parasite associated verminous arteritis (Nielsen et al., 2016). This species has been successfully controlled by anthelmintics, despite recent reports of putative re-emergence associated with reduced frequency of anthelmintic treatments (Nielsen et al., 2012; Nielsen et al., 2016). On the contrary, cyathostomins have become a growing concern in the field (Matthews, 2014; Peregrine et al., 2014). This group of nematodes encompasses 50 known species (Lichtenfels et al., 2008) that infect both young and adult horses (Corning, 2009) and have a ubiquitous distribution throughout geo-climatic conditions (Sallé and Cabaret, 2015a). Their life cycle is direct and usually involves encystment of infective larvae into the caeco-colic mucosa of their hosts (Corning, 2009). In heavily infected horses, *en-mass* emergence of these encysted larvae can cause severe clinical pathology characterized by a loss of weight, colic, diarrhoea, protein-losing enteropathy and the eventually death of the animal (Murphy and Love, 1997; Love et al., 1999).

The use of anti-infectious drugs puts pathogen populations under selection pressure that can ultimately lead to the emergence of resistant or multi-drug resistant populations (Kennedy and Read, 2017). Over the past decades, small strongyle populations, like other livestock-infecting parasitic nematodes (Kaplan and Vidyashankar, 2012), have demonstrated a gradual increase in their resistance to available anthelmintics in every part of the world (Matthews, 2014; Peregrine et al., 2014). Under French settings (Traversa et al., 2012) like in other European (Traversa et al., 2007; Traversa et al., 2009; Relf et al., 2014) or American countries (Slocombe and de Gannes, 2006; Lyons et al., 2008; Molento et al., 2008; Canever et al., 2013), resistant strongyle populations have been reported for every available class of anthelmintics, namely benzimidazoles, tetrahydropyrimidines or macrocyclic lactones. Although previous studies have demonstrated how widespread anthelmintics resistance is, critical assessment of associated risk factors and species composition of resistant parasitic populations is still lacking (Nielsen, 2012), thereby preventing the implementation of clear clear field guidelines. There have been a limited number of reports focusing on factors associated with prevalence of strongyle infection in horses in Germany (Fritzen et al., 2010; Hinney et al., 2011a) or the impact of faeces removal on prevalence in the UK (Corbett et al., 2014). Available studies have considered drenching practices (Lendal et al., 1998; O'Meara and Mulcahy, 2002; Lind et al., 2007; Hinney et al., 2011b; Relf et al., 2012) or estimation of anthelmintic efficacy (Relf et al., 2014; Tzelos et al., 2017). But no attempt has been made to reconcile drenching practices and drug efficacy data. As a consequence, a knowledge gap about putative risk factors and their impacts remains (Nielsen, 2012).

The results reported herein have been gathered as part of a large-scale survey involving 688 horses from 39 French horse farms and riding schools. At each location, an anthelmintic efficacy test and a questionnaire interview about their practices were performed. From this data, risk factors associated with reduced anthelmintic efficacy in equine strongyles have been estimated across drugs. The objective of this study was to bring together anthelmintic efficacy testing and risk factor analysis to provide evidence-based guidelines to the field.

## 2. Materials and methods

### 2.1. Farm and riding school sampling

Our study aimed to evaluate drug efficacy for three drug classes and if possible, to have a control group. Therefore, we aimed to build four groups of at least five horses with a minimal faecal egg count (**FEC**) of 150 eggs/g as recommended by the World Association for the Advancement of Veterinary Parasitology, WAAVP guidelines (Coles et al., 1992). To reach this number of infected individuals, bigger stud farms, i.e. with at least 20 producing mares, were pre-selected from the French Horse and Riding Institute (IFCE) database, in Normandy, Loire Valley, Aquitaine and Burgundy.

Two additional criteria were defined to increase the chance of finding horses with sufficient excretion load to undertake FEC reduction test. First, premises with less than 40 horses were discarded as FEC is usually over-dispersed and focusing on fewer individuals would have reduced our chance to build treatment groups. In Aquitaine however, two farms with slightly smaller herd sizes (25 and 31 horses) were enrolled. Second, last anthelmintic treatment should have been performed three months earlier as this corresponds to the minimal post-moxidectin treatment egg reappearance period (Boersema et al., 1998) advertised on product information (Tzelos et al., 2017). Flyers explaining the purpose of the project were then sent to pre-selected farms before a phone call was made to each manager to make sure that their premises fulfilled requested criteria (at least 40 horses not drenched in the last three months) and to confirm their willingness to participate. Nineteen stud farms were enrolled, *i.e.* five in Normandy, four in Loire Valley, four in Burgundy and six in Aquitaine. Approximately half of these farms (n=11) were involved in horse racing (Thorougbreds and Anglo-Arabians, French trotters or Thorougbreds and other than Thoroughbreds), while the remainder produced leisure ponies (n=4) or leisure horses (n=3) or reared dairy mares (n=2). For each of these, matching riding schools located within each stud farm area were subsequently identified and enrolled for anthelmintic efficacy test, with an additional riding school enrolled in Aquitaine, providing a set of 20 riding schools. This set of matched riding schools was used to investigate putative differences between stud farms and riding schools.

### 2.2. Horse sampling and anthelmintic resistance tests

A first round of faecal sampling was made one week before drenching, to select for individuals with a minimum excretion level of 150 eggs per g (epg) as recommended by the WAAVP guidelines (Coles et al., 1992). Faecal material was stored at 4°C before being processed for faecal egg counting within 24h. Based on individual FEC measured, three treatment groups, *i.e.* fenbendazole (Panacur Equine Guard^®^, Intervet, France), pyrantel (Strongid^®^, Zoetis, France), or ivermectin (Eqvalan pâte, Merial, France), balanced for FEC were built. To do so, individuals were sorted according to their FEC and sequentially allocated to each treatment group. A control group was built in every farm where additional horses with minimal excretion level were available. On day 0, each horse was weighed using a girth tape and orally administered an anthelmintic dose following manufacturer’s requirements. Treatment was administered by the veterinarians enrolled as part of this project.

Faecal material was subsequently taken from each horse 14 days after treatment. Every ivermectin-treated individual still present on the premise 30 days after drenching (n=157 out of 159) was sampled again to identify cases of shortened egg-reappearance period. This short time interval was chosen to minimise disturbance with activities on the premises (horse sales or movements).

### 2.3. Processing of faecal material

FEC were measured by sampling 5g of faecal material for each individual horse, subsequently diluted and thoroughly mixed into 75 mL of a NaCl solution (density of 1.18). Prepared solution was loaded on a McMaster slide and strongyle eggs were counted with a sensitivity of 15 epg.

### 2.4. Questionnaire survey and variable definition

A questionnaire, built upon previous published surveys (Fritzen et al., 2010; Maddox et al., 2012), was used to interview each manager as part of a larger survey on antibiotic and anthelmintic resistance. The anthelmintic-associated questions fell into four categories that addressed global premise overview, available pasture and management, horse health management, and drenching strategy.

For statistical inference, a few variable levels were redefined to avoid redundancy and to provide the analysis with more statistical power. Therefore, one farm that did not apply systematic drenching upon horse arrival was considered as not drenching any horse upon arrival. Rotation between pastures was recoded as occurring either never, or more (frequent) or less (rare) than every 3 months.

In addition, pasture strategies either involved own private pastures dedicated to horses or alternative strategies that included co-grazing with cattle, or access to pastures shared between several breeders. Stocking density was also computed as the herd size divided by available pasture surface, and binned into three categories (more than 5 horses/ha, between 2 to 5 horses/ha or less than 1 horse/ha). A three-level workload variable was defined as the number of horses per worker, falling in either less than 10 horses/employee, between 10 and 20 horses/employee or more than 20 horses/employee.

Two types of managers were defined; those who tried to manage health problems themselves before calling their practitioner and those who called as soon as possible.

Anthelmintic provider was considered as a two-level factor contrasting cases where veterinarians delivered the drug or not. The reasons guiding drenching programs were the same across farms, *i.e.* driven by horse well-being and growth, and were not included further in the risk factor analysis. In the end a set of 21 variables was used (supplementary table 1).

### 2.5. Statistical analyses

#### 2.5.1. Egg Reduction Rate (FECR) and bootstrapping procedure

Sample size to estimate reduced FECR prevalence was determined using EpiTools webserver (Humphry et al., 2004).

When a control group was available, FECR values was computed at farm level by averaging treatment group FEC following (Dash et al., 1988):

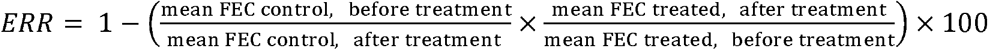

hereafter referred to as **Method 1**.

Otherwise, FECR has been computed following (Coles et al., 1992):

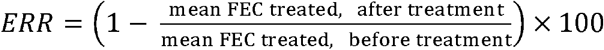

hereafter denoted **Method 2**.

Drug class FECR confidence intervals at the farm level were not estimated as too few individuals were available within each treatment group preventing robust inference of estimate variability (Chernick, 1999). However, for both methods, associated 95% percentile confidence intervals was determined for the region, premise type and drug class and their respective intersections following a block bootstrap approach. This approach takes into account the correlation among observations from the same individual (before and after treatment). For both FECR computation methods, blocks of FECs from the same horse were sampled with replacement from the observed data collected before and after treatment (within region, premise type and drug class), and were used to compute an FECR estimate using equation (1) or (2) accordingly. In that case of method 1, the time-matched control group is used to account for variation in FEC between the two sampling time-points independent of treatment. Therefore, blocks of individual FEC before and after treatment were sampled with replacement from horses belonging to the treated or the control group within a farm.

In both cases, computation was performed 10,000 times to yield the empirical distribution of the FECR from which 2.5 and 97.5% percentiles were sampled to derive the 95% confidence interval.

Pearson’s correlations between FECR of the two methods were estimated using observations from the 24 premises where a control group was available using the *rcorr()* function of the Hmisc package v4.0-3 (Harrell and Dupont, 2017).

#### 2.5.2. Variable selection procedure

To overcome model convergence issues, variables showing too little or no variation across sites (control serology or coproscopy upon horse arrival, faeces removal, access to pasture, horse weight estimation, veterinarian specialization, number of veterinary practices considered for diagnostic) were discarded. Additionally, the total number of veterinary drugs found on-site or the frequency of health register updates were not considered further as determinants of anthelmintic efficacy.

The aim of this study was to quantify risk factors associated with reduced anthelmintic efficacy, hence associating measured drug efficacy with management practices. This requires the selection of variables that would maximize the total variance explained, while avoiding biased estimates through collinearity (Zuur et al., 2010).

Pair-wise Pearson’s correlations between variables were estimated to identify collinearity and avoid redundancy in subsequent modelling of anthelmintic efficacy (supplementary data 2). Any correlation coefficient equal to or above 0.4 was considered as indicative of collinearity. As a result, veterinary advice, health management and the off-licence use of anthelmintics that were accounted for by other variables, were not considered any further.

#### 2.5.3. Dimensionality reduction by Multiple Correspondance Analysis (MCA)

While the discarding of strongly correlated variables addressed the most problematic collinear relationships, significant correlations were still present between the 18 remaining variables (supplementary data 2).

Remaining variables were related to the use of anthelmintics, the use of pasture, or the general constraints applying to the premise (work load measured by the number of horses per employee, and annual veterinary expenses per horse). For each of these three categories, a multiple correspondence analysis was performed on the dedicated set of variables with the FactoMineR v.136 package (Lê et al., 2008). Premise coordinates on the first two components of the MCA were subsequently used to define three-level (premise constraints variable) or four-level variables (anthelmintics and pasture usage).

#### 2.5.4. Marginal modelling of drug efficacy

A marginal modelling approach of individual horse egg count reduction rate (FECR) was applied as outlined elsewhere (Walker et al., 2014) and implemented in R (R Core Team, 2016) with the geepack package v1.2-1 (Højsgaard S., 2006). In that framework, individual egg counts measured at a given time, before or after treatment, are assumed to be Poisson distributed and thus modelled with a log-linear regression model.

This model includes environmental variables (premise type, region, and the three variables built from the MCA) interacting with a binary variable coding for the treatment, *i.e.* taking value of 1 after treatment or 0 before treatment. This variable accounts for the treatment-mediated change in egg count reduction, hence estimating FECRs, while the fitted interactions estimate the contribution of considered environmental conditions to FECRs (Crellen et al., 2016). Exponentiated estimates therefore provide the relative risk of increased (relative risk above one)/decreased (relative risk smaller than one) FECRs associated with a given environmental variable. Any variable whose relative risk confidence interval does not include one is declared as significantly impacting on the FECRs. A forward-backward procedure was implemented with the *stepAIC()* function of the R MASS package v7.3-47 to select for the model minimizing the Akaike Information Criterion.

The model aimed to quantify universal risk factors, *i.e.* considered environmental effects across drug class. Treatment group was thus added into the model. Drug class-specific analyses based on individual treatment group data taken separately did not provide reliable results and were not reported.

## 3. Results

### 3.1. Observations from questionnaire surveys

Detailed answers from questionnaire surveys are provided as supplementary data 1.

#### 3.1.1. Overview of the enrolled premises

At least three treatment groups could be built in every location. However, herd size was highly variable between sites, ranging from 21 to 250 individuals (mean herd size of 70 horses). This variation was mostly driven by the herd being bigger in stud farms than in riding schools (average herd size of 88.3 and 52.2 respectively; p=0.008). Horses were generally housed in individual boxes (n=31/39) and a few premises had an outdoor-only breeding system (n=7/39). Noticeably, staff size did not strongly correlate with herd size (r=0.33, p=0.04), especially in stud farms where workers were in charge of 13 horses more than in riding schools (p<10^−4^).

#### 3.1.2. Horse movements on site

Horse movement occurred in half of the premises (n=21/39) at least once a month while seven premises were rarely housing horses from other places. For these introductions, no serology, no coproscopy and no anthelmintic efficacy tests were performed in any of the 39 premises, whereas anthelmintic drenching upon arrival was implemented in 11 riding schools and seven stud farms. Only four managers reported seeking advice from their veterinarians to manage these movements.

#### 3.1.3. Pasture availability and management

In every site, horses had outdoor access and could generally grazed throughout the year (n=27/39). Most premises had their own pastures that were grazed by horses only. Three farms had access to pastures shared with other breeders, and mixed grazing of horses with cattle was implemented in seven premises. Stocking density was below 3 horses/ha in 75% of sites. However, it was as high as 14 horses/ha and was generally higher in riding schools than in stud farms (average densities of 1.3 and 4.3 horses/ha respectively, p=2x10^−4^). Rotation between pastures was implemented in 29 locations at least once a year and driven by grass growth. Faeces removal was implemented in one premise. Manure spreading was performed in one third (n=10/39) of the surveyed sites.

#### 3.1.4. Health management and interactions with veterinarians

About two-thirds of the premises relied on specialized equine practitioners (n=24/39), who were often called after managers had already attempted to manage health problem themselves (n=28/39). Half of the premises (n=20/39) were consulting several practices to cross-validate advice, or benefit from several skills, or both.

Yearly veterinary expenses per horse varied from less than 100 € (n=15), between 100 and 200 € (n=14), or more than 200 € (n=10).

Importantly, managers reporting to be more independent in health management were over-represented in sites not implementing any measure upon horse introduction (13/39) and in the sites spending less than 100€/horse/year (*r*=−0.39 and −0.41 respectively; supplementary table 1). Health management was hence confounded by the two other variables and not considered further.

Mandatory on-site health register was variably used, *i.e.* 20 managers fulfilled it regularly (systematically or on a regular basis), while 19 rarely did it (never or doing it from time to time). The number of veterinary drugs found within on-site pharmacy greatly varied from null to 15, with two-thirds of premises having 5 drugs or less with a slight trend of more medications found in horse riding schools (Kruskal-Wallis test, p=0.07).

#### 3.1.5. Drenching strategy for intestinal nematodes

Anthelmintic dosing was usually based on a visual weight estimate (n=27/39) that could be combined with girth tape (n=9/39), and rarely with a scale (n=2/39). Grouped-based drenching was carried out in 11 sites. Time of drenching was registered most of the time (n=31). Drenching frequency occurred two (n=13), three (n=11) or four times (n=14) a year, and drenching programs alternated between drug classes.

Noticeably, a limited fraction of premises (n=6) reported off-license use of anthelmintics despite most of the managers seeking advice from their veterinarians (5 out 6). These involved ivermectin (n=3), doramectin (n=2) or praziquantel (n=2) licensed for ruminants.

The off-licence use of anthelmintics was only reported in premises implementing two (n=5) or three (n=1) annual treatments, resulting in a negative correlation between these two variables (r=−0.41, supplementary data 2). The off-licence variable was hence not considered further in the modelling analysis.

Anthelmintics were bought from veterinarians in 62% of cases, while three and 16 managers reported buying from the internet or their pharmacist, respectively. In the latter case, only three managers were aware of the legal requirement of showing a veterinarian’s prescription to the pharmacist.

The delivery of anthelmintics by veterinarians occurred more frequently when they contributed to the drenching scheme design, *i.e.* 79 % of cases against 40% when they did not (*r*=0.4, supplementary table 2). In addition, stud farm managers relied more on their veterinarian’s advice for drenching in comparison to their riding school counterparts (84% of stud farms vs. 67 % of riding schools; *r*=0.45, supplementary table 2). Premises that did not rely on veterinary advice also did not implement any measure upon horse arrival (*r*=0.47, p=0.003; supplementary table 2). Veterinary advice was hence accounted for three other variables, and was not considered further in the analysis.

FEC-based drenching regimen was implemented in 14 premises.

### 3.2. Results of anthelmintic efficacy tests

This design provided enough resolution to detect prevalence as low as 1%, with precision of 0.05 and assuming FEC sensitivity of 70% and specificity of 90%.

A total of 688 horses from 39 premises were sampled at least once during this experiment. Out of these, 601 horses excreting more than 150 epg before treatment were enrolled for the anthelmintic resistance test (Table 1). Control groups were available in 24 out of the 39 retained farms (Table 1). Average FEC before treatment was 912 ± 762 epg (supplementary data 3).

**Table 1.**
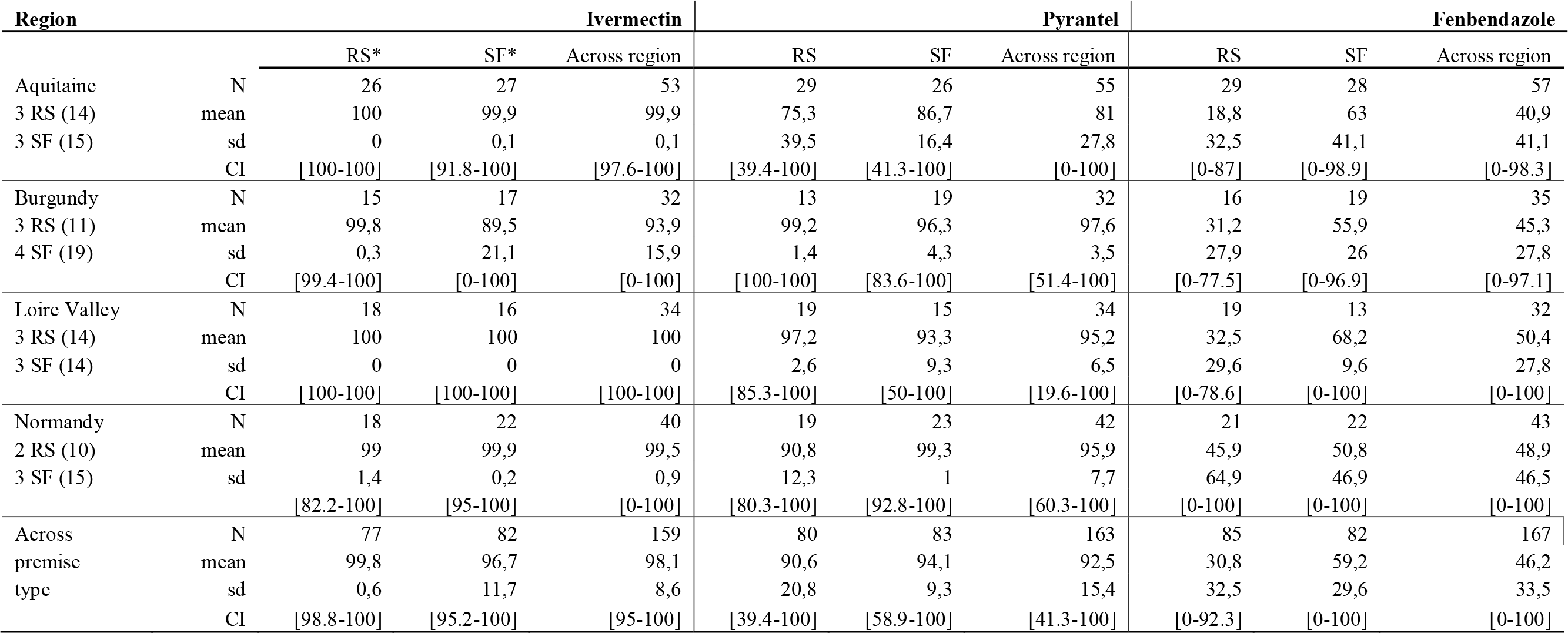
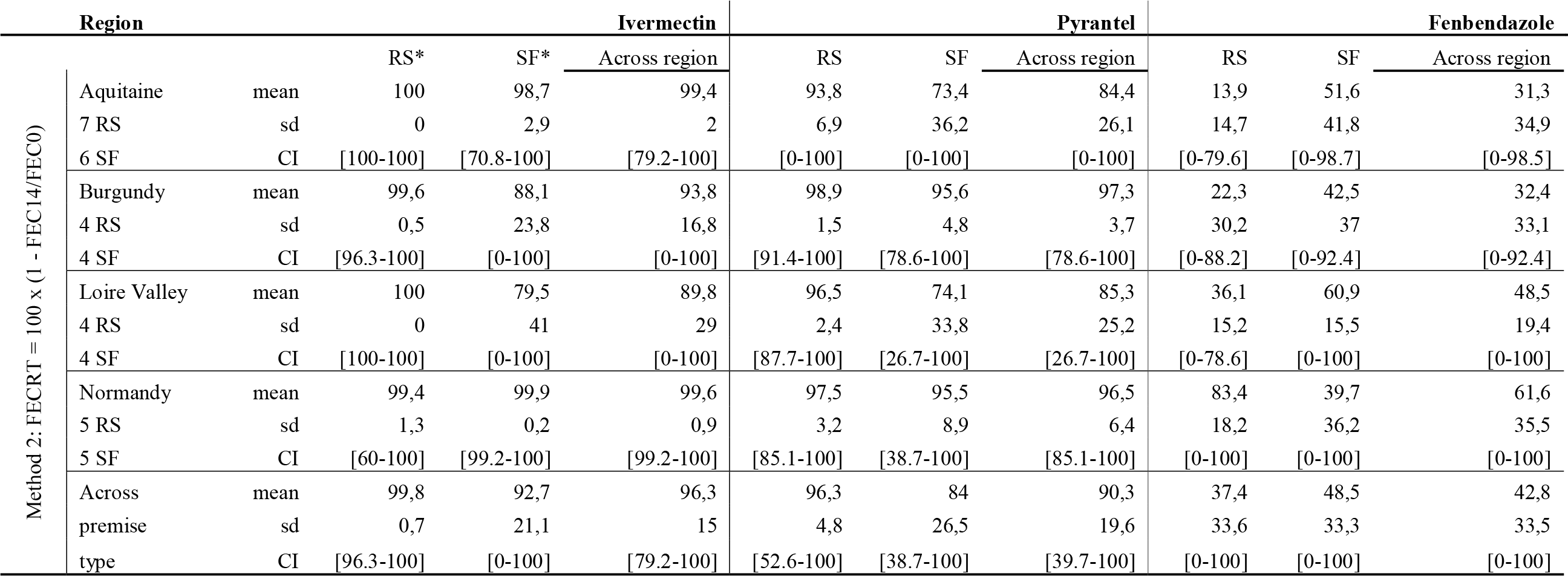
Average Egg Reduction Rate estimated across premise type and regions. Drug-specific average Egg Reduction Rates (mean) and standard deviations (sd) measured 14 days after treatment have been collated for each drug and region of interest for the two egg reduction rate calculation methods used. CI stands for Cross-sectional confidence intervals. N indicates the number of horses available, while RS and SF stand for riding-school and stud farm respectively. Figures in brackets under the Region column stands for the number of horses allocated to the control group in Riding Schools or Stud Farms accordingly.

Estimated Faecal Egg Count Reduction (FECR) and associated variation have been reported in Table 1 while premise-level estimated FECR have been attached as supplementary data 4. FECRs measured in the 24 sites where control group was available showed highly consistent results between the two implemented calculation methods for ivermectin and fenbendazole (Pearson’s correlation coefficient of 100 and 82%, respectively). This correlation however dropped to 65% for pyrantel.

Estimated FECRs demonstrated the almost generalized inefficacy of fenbendazole with an average FECR of 46.2% (sd=33.5%) or 42.8% (sd = 33.4%) for method 1 and 2, respectively and confidence intervals not including 100% efficacy in Burgundy and Aquitaine (Table 1, supplementary data 4). Nevertheless, two riding schools and one stud farm located in Normandy exhibited FECRs of at least 90% (supplementary data 4).

Observed trends for ivermectin were the exact opposite of these, as the mean estimated FECRs were 98.1% (sd: 8.6%) and 96.3% (sd: 14.5%) according to methods 1 and 2, respectively (Table 1, supplementary data 4). Seven horses from three riding schools and three stud farms exhibited FECRs lower than 90% after ivermectin treatment, resulting in bigger confidence intervals in Aquitaine and Normandy (Table 1). Egg reappearance was investigated 30 days after ivermectin drenching, *i.e.* based on FEC from 157 available horses. At this time, only nine horses excreted strongyle eggs (mean FEC of 14.6 epg). These were found in eight farms (three from Burgundy, four from Aquitaine and one from Loire Valley) and one riding school from Aquitaine. Only three of these horses had egg excretion levels above 50% of their before treatment FEC.

Pyrantel exhibited an intermediate profile in comparison to the two other drugs as average FECRs were close to the 90% threshold, *i.e.* 92.5% (sd: 15.4%) and 90.3% (sd: 19.6%) for methods 1 and 2, respectively (Table 1).

### 3.3. Risk factors associated with anthelmintic efficacy

The variation in FEC before and after treatment was explicitly modelled by considering every individual FEC and by correcting for the farm environmental variables. This is to better capture the inter-individual variations associated with FEC while estimating the relative risk associated with environmental variables. Relative risks associated with reduced FECR were estimated across drug categories, any relative risk above 1 indicating an increased egg count after treatment and thus reduced efficacy. Drug-specific risk factors were subsequently estimated considering observations from each treatment group independently.

#### 3.3.1. Multivariate analysis and summary variables

An MCA was applied to the set of variables related to the use of pasture and anthelmintics (figure 1) or the constraints applying to the premise (figure 2) to avoid fitting collinear variables in the drug efficacy model.

**Figure 1.**
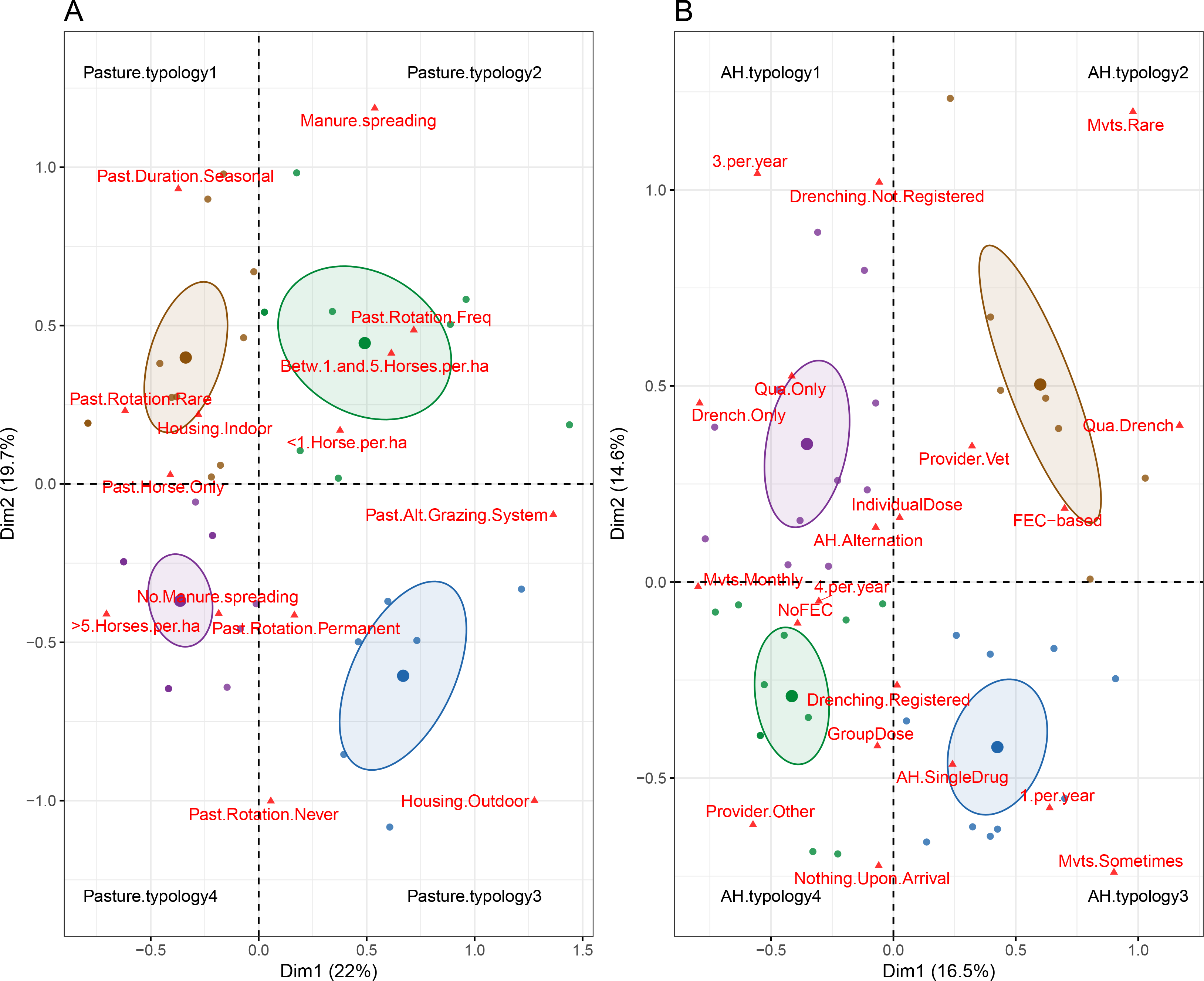
Multiple correspondence analysis of the variables related to pasture (A) and anthelmintics (B) uses. The first two components of the analysis are plotted and distinguish between four different typologies annotated in black (e.g. pasture.use1). Environmental variables are represented by red triangles while dots represent corresponding premises, colored according to the typology they belong to. Ellipses represent the 95% credible interval associated with each typology.

**Figure 2.**
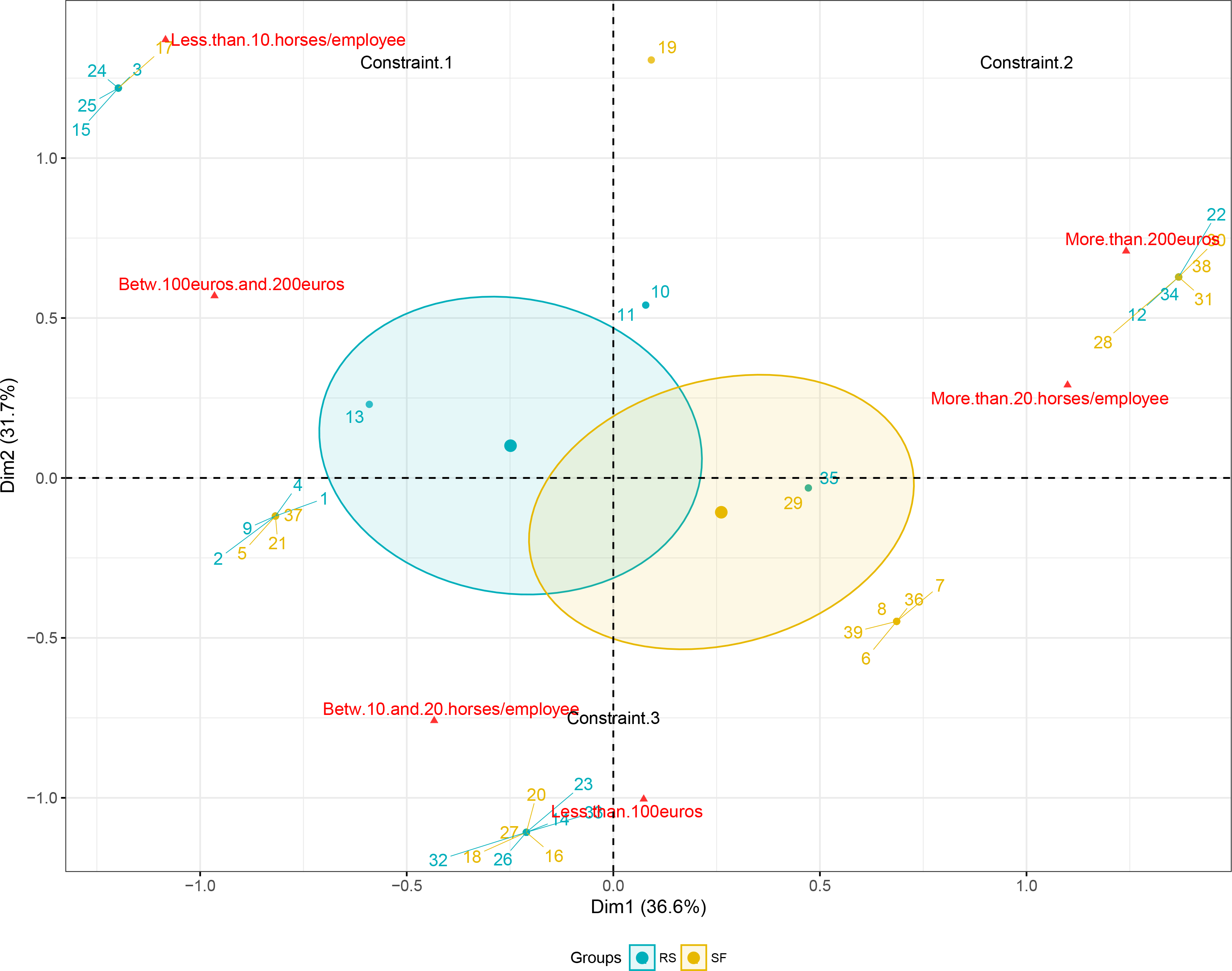
Multiple correspondence analysis of the variables related to structural constraints applying to premises. The first two components of the analysis are plotted and distinguish between three different typologies annotated in black (e.g. Constraint.1). Levels corresponding to the considered workload and veterinary expenses variables are represented by red triangles while dots represent corresponding premises, colored according to the typology they belong to. Ellipses represent the 95% credible interval associated with each typology.

The MCA applied to the variables related to pasture usage accounted for 42% of the between-variable variance (figure 1A) and distinguished between four contrasted typologies. The first typology regrouped premises where grazing options were limited, *i.e.* mostly indoor housing and seasonal grazing with rare rotation between pastures. While typologies 1 and 3 were mostly driven by the housing system, the split between the other two systems was accounted for by the pasture surface availability (figure 1A). The 2^nd^ pasture usage typology clustered together premises with the lowest stocking density, that were also able to perform manure spreading and frequent rotation between pastures. These four typologies were used to build a pasture usage variable for subsequent modelling.

The same approach resolved 31% of the variation in anthelmintics usage between premises (figure 1B). In this case, typologies 2 and 4 distinguished between premises with the most extreme behaviors regarding drug use. Premises falling under typology 2 applying strategies usually regarded as the most sustainable for drug resistance and typology 1 represented an intermediate situation between typologies 2 and 4. Typology 3 grouped together premises relying on a single annual macrocyclic lactone treatment. These four typologies were used to build an anthelmintic usage variable for subsequent modelling.

The two remaining variables to be considered addressed the annual veterinary expenses and the workload in premises (figure 2). These two variable were summarized as a so-called “constraint” variable that distinguished between three situations. Typology 2 described the most heavily constrained premises with highest workload and veterinary expenses (figure 2), whereas typology1 and 3 accounted for the smallest workload and most reduced veterinary expenses respectively (figure 2). Typology 1, defined by more limited workload and intermediate veterinary expenses, mostly encompassed riding schools. However, he correlation between the premise type and the three constraint levels was not significant (*r*=0.26, *p*=0.1, supplementary table 5).

No residual significant correlations existed between variables considered for modeling (supplementary table 5).

#### 3.3.2. Risk-factors across anthelmintic drug class

Modeling of drug efficacy relied on a set of six variables, *i.e.* day of treatment, treatment group, region and the three summary variables derived from MCA.

A first analysis investigated universal factors associated with drug efficacy, measured by FECR, that would be true across anthelmintic drugs and would not depend on the drug mode of action. Relative risks associated with the retained variables have been provided in table 2. Noticeably, the interaction term between the day of treatment and premise type or the region of interest or premise constraints were not retained by the model selection procedure, suggesting these variables were not providing information to the modelling of drug efficacy that had not been already accounted for by other variables.

**Table 2.** Estimated odd ratios associated with retained environmental variables. For each of the retained management practice, the relative risk of reduced (below 1) or increased (higher than 1) risk of anthelmintic resistance is provided. Associated 95% confidence intervals lower and upper limits are given as well as the associated *p* value.

| Variable | Odd ratio | Lower Odd Ratio | Upper Odd Ratio | *p* |
| --- | --- | --- | --- | --- |
| Ivermectin vs. Fenbendazole | 0,02 | 0,01 | 0,08 | <10<sup>-2</sup> |
| Pyrantel vs. Fenbendazole | 0,14 | 0,08 | 0,22 | <10<sup>-2</sup> |
| Pasture.use2 vs Pasture.use1 | 0,53 | 0,36 | 0,79 | <10<sup>-2</sup> |
| Pasture.use3 vs Pasture.use1 | 0,78 | 0,52 | 1,18 | 0,24 |
| Pasture.use4 vs Pasture.use1 | 0,78 | 0,55 | 1,11 | 0,17 |
| AH.use2 vs AH.use1 | 0,57 | 0,36 | 0,92 | 0,02 |
| AH.use3 vs AH.use1 | 0,89 | 0,62 | 1,27 | 0,52 |
| AH.use4 vs AH.use1 | 1,10 | 0,78 | 1,57 | 0,58 |

In line with estimated FECRs, pyrantel and ivermectin were less at risk of reduced FECRs than fenbendazole considered as the reference level (Table 2).

Premises falling under the second typology of pasture usage (figure 1A) demonstrated a significant reduction of drug resistance risk (OR=0.53, p=0.001) in comparisons to other modalities (Table 2). This typology equally matched riding schools (n=6) or stud farms (n=5) but was over-represented in Normandy (6/11). This certainly explains why the site location was not retained in the model selection procedure.

The second typology of anthelmintic usage (figure 1B), that regroup strategies usually thought of as more sustainable toward drug resistance, was significantly associated with a reduced risk of drug resistance (OR=0.57, p=0.02). No significant differences could be made between other typologies (Table 1).

## 4. Discussion

Current knowledge about anthelmintic resistance in equine strongyles is usually scattered across drug efficacy reports and questionnaire surveys about parasite management (Nielsen, 2012). This leaves a major knowledge gap in the critical assessment of factors underpinning anthelmintic resistance in equids. Our study aims to fill in this gap with the report of an association between anthelmintic FECRs and management practices in horses.

The drug efficacy landscape in the present study remained similar to what was reported in a previous study in France (Traversa et al., 2012) and what was described in other countries (Matthews, 2014). In summary, fenbendazole cannot be used for the management of small strongyles any more, in contrast to ivermectin whose efficacy remained above 95%. Pyrantel had an intermediate efficacy FECR pattern with a 90% reduction of egg-excreting animals. However, two original findings tend to depart from this general pattern. First, the risk of fenbendazole resistance was significantly lowered in Normandy with a few premises (3/39) still harbouring fenbendazole-susceptible strongyle populations. This was in line with a previous study (1/18) conducted in France (Traversa et al., 2012). However, it was not possible to identify an obvious consistent factor that would explain this sustained fenbendazole efficacy. In-depth investigation of practices and analysis of parasitic community structure with a nemabiome approach (Avramenko et al., 2015) may help better understanding this feature and then confirming fenbendazole-susceptibility by interrogating beta-tubulin sequences and allelic frequencies (Lake et al., 2009). Second, our results showed that ivermectin efficacy may not be sustained at its current level in the near future as egg excretion already took place 30 days after treatment in a limited number of horses across all regions. In addition, larger FECR confidence intervals were encountered in Normandy and Aquitaine, suggestive of a higher variability in ivermectin efficacy. Original ERP was 9 weeks for ivermectin (Boersema et al., 1996) but indications of shortened ERP have been found in Germany (von Samson-Himmelstjerna et al., 2007), the UK (Daniels and Proudman, 2016), Belgium, the Netherlands and Italy (Geurden et al., 2014) and had never been reported in France. In this study, the quantitative determination of ERP was not possible due to practical reasons. Instead we focused on the 30^th^ day post ivermectin treatment to minimize interferences of our design with activities on premises and to ensure that most of the treated horses would still be available for sampling. Despite this, a few horses had already been sold or sent to other premises for training.

Beyond the crude estimation of drug efficacy, this study aimed to identify major determinants underpinning egg reduction rate, and to estimate their respective relative contributions to provide evidence-based recommendations in the field.

Pasture-related variables were significant contributors to the variation in drug efficacy measured by egg reduction rate. Noticeably, sites with typology 2, that had the lowest stocking density, were able to implement frequent pasture rotation and to perform manure spreading, had a significantly decreased risk of drug resistance. Stocking density has been advocated as a factor driving drug resistance even if it could not be associated with more elevated infection rate in a German epidemiological study (Fritzen et al., 2010). On the contrary, frequent pasture rotation is one of the evasion strategy recommended to minimize pasture contamination (Michel, 1985). Therefore, it is probable that the combination of both a reduced horse density with frequent rotation between pastures minimized the use of drugs in these sites. The effect of manure spreading in the context of drug resistance remains uncertain. It could simply correlate with the available grazing surface and mirror the stocking density but this correlation was not significant in this dataset (*r*=−0.28, *p*=0.09).

Interestingly, drug resistance was significantly reduced in sites implementing a combination of FEC-based drenching programs, determination of drug dosage on an individual basis and a high level of biosecurity, *i.e.* little horse movement combined with a qurarantine and drenching upon horse introduction. Evidence-based drenching and individual drug dosage have long been advocated for in ruminant and equine systems as a sustainable parasite management practices, as the former is thought to reduce selection pressure (Kenyon et al., 2009) and the latter is thought to prevent under dosing whose impact on drug resistance development relies on many parameters (Smith et al., 1999; Silvestre et al., 2001). Our findings thus provided evidence to promote their enforcement in the field.

Notably, the more sustainable anthelmintic usage typology (typology 2) also accounted for the delivery of anthelmintic treatment by veterinarians. Under French regulations, veterinary medications can be delivered by veterinarians or pharmacists upon display of a veterinary prescription (Anonymous, 2007). This regulation was generally applied across the considered study sites and well correlated with the involvement of veterinarians into the drenching scheme design. Recent study in the UK suggests that practitioners may provide useful advice on drug use (Easton et al., 2016) and thus reinforce their role in sustainable parasite control. Such partnerships between veterinarians and horse owners should thus be encouraged in France as 40% of the considered sites designed their own drenching scheme.

These first insights into determinants of drug efficacy only focused on environmental factors, putting aside intrinsic worm characteristics, like species composition, that should be investigated further. Recent advances in parasite metagenomics would help addressing this question.

### Conclusions

This study reports the first risk estimation analysis between management practices and drug efficacy in equine strongyles. While drug resistance prevalence remains in agreement with previous surveys from France and other countries, *i.e.* a generalized failure of fenbendazole, a decreasing efficacy of pyrantel and reasonably high efficacy of ivermectin despite evidence of reduced egg reappearance period. Most importantly, we have quantified the relative risks and benefits associated with equine farms management practices. These estimations provided support to advocate for FEC-based treatment regimens as well as individual determination of anthelmintic dosage. In addition, tight biosecurity enforced by reduced horse movements and a combinaison of anthelmintic treatment with quarantine upon horse introduction should be recommended. Anthelmintics delivery by veterinarians was also among beneficial factors relative to drug resistance. Also, sites with frequent pasture rotations and stocking rate below five horses/ha displayed a reduced risk of drug resistance.

---

**Supplementary Data 1. Retained variables and data distribution across premises.**

**Supplementary Data 2. Pair-wise Pearson’s correlations between variables**

**Supplementary Data 3. Average faecal egg count by premise type and region before anthelmintic treatment**

**Supplementary Data 4. Farm-level egg reduction rates with associated confidence intervals**

**Supplementary Data 5. Correlation between variables retained for the modelling of Faecal Egg Count Reduction**

## Acknowledgements

Authors are greatly indebted to the premise managers who took part to this study and gave us access to their stud or riding schools. We would like to acknowledge Pr. C. Chartier for kindly granting access to his parasitology laboratory, Dr. C. Laugier for discussing the questionnaire and Dr. L. Crespin for critical comments on this manuscript. We also would like to acknowledge the technical support of Dr. C. Charvet, C. Koch and C. Musset during this survey, and J. Noonan, A. Tracey and P. Driguez for their review of the English language. This work has been funded by the French Institute for Horse and Horse Riding (IFCE) and the Fonds Eperon fund for horse racing (BIOREQUI project), as well as the Merial and Zoetis companies.

